# Density-dependent environments can select for extremes of body size

**DOI:** 10.1101/2022.02.17.480952

**Authors:** Tim Coulson, Anja Felmy, Gioele Passoni, Robert A Montgomery, Jean-Michel Gaillard, Peter HJ udson, Joseph Travis, Ronald D Bassar, Shripad Tuljapurkar, Dustin J Marshall, Sonya Clegg

**Affiliations:** Department of Biology, University of Oxford, Oxford, OX1 3SZ, UK; Department of Evolutionary Biology and Environmental Studies, University of Zurich, Switzerland; Department of Biological Science, Florida State University, Tallahassee FL 32306, USA; Laboratoire de Biometrie et Biology Evolutive, University of Lyon 1, Lyon, France; The Huck Institutes, Penn State University, State College, PA 16802, USA; Department of Biology, Williams College, Williamstown, MA 01267, USA; Department of Biology, Stanford University, Palo Alto, CA 94305, USA; School of Biological Sciences, Monash University, Melbourne, Victoria, Australia 3800

**Keywords:** Carrying capacity, Dwarfism, Gigantism, Integral Projection Model, Life history evolution

## Abstract

Body size variation is an enigma. We do not understand why species achieve the sizes they do, and this means we also do not understand the circumstances under which gigantism or dwarfism is selected. We develop size-structured integral projection models to explore evolution of body size and life history speed. We make few assumptions and keep models simple: all functions remain constant across models except for the one that describes development of body size with age. We set sexual maturity to occur when size attains 80% of the asymptotic size, which is typical of a large mammal, and allow negative density dependence to only affect either reproduction or juvenile survival. Fitness – the quantity that is maximized by adaptive evolution – is carrying capacity in our models, and we are consequently interested in how it changes with size at sexual maturity, and how this association varies with development rate. The simple models generate complex dynamics while providing insight into the circumstances when extremes of body size evolve. The direction of selection leading to either gigantism or dwarfism crucially depends on the proportion of the population that is sexually mature, which in turn depends on how the development function determines the survivorship schedule. The developmental trajectories consequently interact with size-specific survival or reproductive rates to determine the best life history and the optimal body size emerges from that interaction. These dynamics result in trade-offs between different components of the life history, with the form of the trade-off that emerges depending upon where in the life history density dependence operates most strongly. Empirical application of the approach we develop has potential to help explain the enigma of body size variation across the tree of life.

## Introduction

Body size evolution, particularly when resulting in either dwarfism or gigantism, has long fascinated biologists. Stout infantfish (*Schindleria brevipinguis*) achieve sexual maturity at less than 0.1g (Watson and Walker, 2004) while blue whales (*Balaenoptera musculus*) can grow to weigh 150 tonnes representing a span in adult weights of over nine orders of magnitude. Lifespan in vertebrates is not quite so variable, but the range of three orders of magnitude is still impressive: Greenland sharks (*Somniosus microcephalus*) can live up to half a millennium (Nielsen et al., 2016), while the coral reef fish, the seven-figure pygmy goby (*Eviota sigillata*), is elderly if it survives for two months (Depczynski and Bellwood, 2005). There are physiological limits that define the extremes of body size and longevity in vertebrates (Goldbogen, 2018), but the selective forces that may push organisms towards these extremes are presently unclear.

Body size variation across species is statistically associated with life history variation in an allometric manner (Savage et al., 2004; West et al., 1997). As size increases, there is also an increase in the value of traits measured in units of mass (e.g. neonatal mass), time (e.g. life expectancy), and length (e.g. body length). In contrast, as size increases, the values of traits describing the frequency of events, such as reproductive rates, decrease (Savage et al., 2004; West et al., 1997).

Within species, patterns of size variation are less clear. While body size has very often been found to be under directional selection, it has rarely been found to evolve in line with predictions (Kingsolver et al., 2001; Merilä et al., 2001). Body size evolution remains challenging to understand because identical processes can result in increases in body size and a slowing of the life history in some species, yet the exact opposite in others: food limitation selects for an increase in body size at sexual maturity in some species of fish (Travis et al., 2014), but a decrease in ungulates (Ozgul et al., 2009; Raia and Meiri, 2006). We do not have a good understanding of why species are the size they are (Audzijonyte et al., 2020).

Darwinian demons are hypothetical creatures capable of simultaneously maximizing all components of fitness (Law, 1979). In doing so, they achieve sexual maturity immediately after birth, continuously produce litter sizes of an infinite number of viable young, and are immortal. They would presumably be tiny, perhaps infinitesimally so, given development takes time. Regardless of their size, we would instantly be neck-deep in such pests. Fortunately, Darwinian demons do not exist because all individuals face trade-offs.

Trade-offs occur when something prevents all components of fitness being maximized simultaneously (Kozłowski et al., 2020; Stearns, 1992; Stearns, 1977). They arise at the individual level when something limits population growth. In the absence of trade-offs, populations grow exponentially and organisms evolve towards a Darwinian demon life history as allocation of resources to early reproduction is always favored under such circumstances (Coulson, T Benton, et al., 2006; McGraw and Caswell, 1996). Energy availability is frequently assumed to be the constraint that generates trade-offs (B Kooijman and S Kooijman, 2009), but the availability of enemy-free space, breeding sites, water, or other molecules essential for life, can also generate them. The question we are interested in is how trade-offs can select for long developmental periods, large body size, and slow life histories, the apparent antithesis of Darwinian demons?

Approaches to understanding both life history and body size evolution often involve specifying a limiting factor and a life history trade-off, before identifying the fittest strategy. For example, in bioenergetic and dynamic energy budget models, energy is assumed to be limiting, the trade-off is specified via rules determining the allocation of energy to maintenance, development, and reproduction (B Kooijman and S Kooijman, 2009), and the fittest strategy is identified usually via an evolutionary game (Day and Taylor, 1997; Koziowski and Weiner, 1997; Kozłowski, 1992). A related approach involves agnosticism as to the limiting factor, *a priori* specification of a trade-off between two components of the life history such as offspring number and offspring size (Smith and Fretwell, 1974), and use of an evolutionary game to identify the fittest strategy (Childs et al., 2004; Grant, 1997; Meszena et al., 2002; Metcalf et al., 2008; Roff, 1993). A third alternative is to identify the quantity that evolution maximizes (e.g., fitness), and to examine how independently altering each part of the life history impacts fitness. Selection is then assumed to predominantly operate via the life history components with the largest sensitivities of fitness (Caswell, 2001; Holland-Jones and Tuljapurkar, 2015; Tuljapurkar et al., 2009). This approach has been used for deterministic, density-independent environments where fitness is the population growth rate measured as λ, and stochastic, density-independent environments where fitness is the long-run stochastic growth rate (Tuljapurkar et al., 2009).

The last approach has the virtue of making few assumptions as it does not require specification of a tradeoff, but it does suffer from a shortcoming in that continuous population growth occurs in the absence of population limitation, and therefore environmentally-determined trade-offs shaping evolutionary trajectories may not exist. We address this limitation by using the approach in negative density-dependent environments where population limitation, by definition, exists (Turchin, 1999). This imposes a constraint on population growth and mean lifetime reproductive success, but does not require us to make any assumptions about the nature of the limiting factor (e.g. energy or enemy-free space), and we do not need to specify a trade-off *a priori.* Instead, the trade-offs emerge as a function of where in the life cycle limiting processes operate most strongly, and where they are absent [see also Charlesworth (1994).

When trade-offs reveal themselves via the imposition of population limitation, population size will achieve an equilibrium referred to as carrying capacity *K*, density dependence will be observed, and the population growth rate will equal zero. It is tempting to equate density dependence with food limitation (White, 2008), but that is too narrow a definition. Density dependence is simply a statistical pattern where no long-term temporal trend in population numbers is observed. Any limiting process can generate density-dependent dynamics (Turchin, 1999). In deterministic, density-dependent environments, regardless of the limiting process, carrying capacity has repeatedly been proven to be fitness, i.e. the quantity maximized by evolution (Charlesworth, 1973,1994; Kentie et al., 2020; Lande, Engen, and Sæther, 2017; Lande, Engen, and Saether, 2009; MacArthur, 1962; Mylius and Diekmann, 1995; Takada and Nakajima, 1992,1998). The strategy that has the highest value of carrying capacity is evolutionarily stable (appendix) and cannot be invaded by any strategy with a lower carrying capacity (Kentie et al., 2020).

We are interested in understanding the evolution of extremes of body size, so we develop size-structured models (that are density-dependent), and we examine how altering growth trajectories impacts body size, life history speed, and carrying capacity while imposing a constraint that sexual maturity occurs at a fixed proportion of asymptotic size. We discover that:

1. The key parameter determining selection on size at sexual maturity and life history speed is the proportion of the population that is sexually mature. The proportion reflects a balance between juvenile survival and adult life expectancy. This result generalizes previous work that did not consider body size but that characterized the role of comparative juvenile and adult survival rates on life history evolution (Charlesworth, 1973; Takada and Nakajima, 1992).
2. Delaying sexual maturity generates a mortality cost to juveniles, such that a smaller proportion of each cohort survives to maturity. If this cost is offset by a survival or reproduction benefit to adults, via either an increase in life expectancy or increased reproduction, then larger body sizes and slower life history strategies will be selected. If the juvenile mortality cost is not offset by the adult fitness benefit, then small body sizes and faster life histories that are closer to that of Darwinian demons will evolve.
3. In our models, carrying capacity is fitness and density dependence generates these trade-offs. In density-dependent environments population growth and mean lifetime reproductive success both equal one at equilibrium. Evolution acts to maximize carrying capacity by suppressing the value of negatively density-dependent demographic rates (here, reproduction in one scenario and juvenile survival in the other). As these rates are suppressed, those that are not density-dependent (which rate depends upon the scenario) will increase in order to maintain a population growth of one.
4. The simultaneous suppression of density-dependent rates and increase in density-independent rates generates the life history trade-offs we observe. Where in the life history these trade-offs occur depends upon which demographic rates are influenced by density, and which are not.
5. The cross-life trade-offs we identify could be generated by density-independent processes such juveniles and adults experiencing different environments as well as by the density dependence on which we focus.

## Methods

### Overview of approach

We develop simple models where only one demographic rate is density-dependent. In scenario 1, reproduction is negatively density-dependent; in scenario 2, juvenile survival is negatively density-dependent. Within each scenario we construct 20 models, each describing a unique life history strategy. These strategies differ from one another in the growth trajectory that individuals follow. The different growth trajectories result in different asymptotic sizes and sizes at sexual maturity across life histories. We can consequently distinguish each life history by its size at sexual maturity. By comparing fitness across strategies within a scenario we can explore selection on life history strategy (Kentie et al., 2020; Tuljapurkar et al., 2009).

In life history theory, evolution maximizes the mean fitness of a strategy (Metcalf et al., 2008; Stearns, 1977). Mean fitness of a life history strategy is always a quantity that describes some aspect of the strategy’s population dynamics (Charlesworth, 1994; McGraw and Caswell, 1996; Tuljapurkar, 1990). In deterministic density-dependent environments where competition between individuals is symmetric – the case that interests us – the quantity that evolution maximizes is well-known to be carrying capacity *K* (Charlesworth, 1973, 1994; Kentie et al., 2020; Lande, Engen, and Sæther, 2017; Lande, Engen, and Saether, 2009; MacArthur, 1962; Mylius and Diekmann, 1995; Takada and Nakajima, 1992, 1998). The life history strategy with the highest carrying capacity will always be evolutionarily stable (Charlesworth, 1994; Kentie et al., 2020). We can consequently identify the evolutionarily stable life history strategy by comparing carrying capacities across different strategies. Our first aim is to understand how evolution maximizes carrying capacity within each scenario, and we do that by identifying the demographic rates that determine the value of K.

Evolution alters the values of demographic rates to maximize carrying capacity via optimizing survivorship and fertility schedules (Kozłowski et al., 2020; Stearns, 1977). Survivorship describes the probability of surviving from birth to each age, while fertility schedules describes the production of offspring at each age. Our second step is to explore how these schedules are optimized to maximize K. By combining these steps we gain insight into the circumstances when extremes of body size are expected to evolve.

We make few assumptions, and strive to keep models simple, while choosing forms of demographic functions that are typical of those observed in nature such as an increase in survival rate with body size, and a juvenile and adult stage either side of sexual maturity. Terms used in the text are defined in Table 1.

**Table 1.** Notation used in the paper. Please see Table 2 for values of model parameters.

| Term | definition |
| --- | --- |
| $a$ | Age |
| $a_m$ | Age at sexual maturity |
| $\beta_0, \alpha_0, \rho_0, \gamma_0$ | Function intercepts (survival, development, reproduction, inheritance) |
| $\beta_z, \alpha_z, \rho_z$ | Function slopes for body size (survival, development, reproduction) |
| $\beta_N, \alpha_N, \rho_N$ | Function slopes for density (survival, development, reproduction) |
| $\alpha_v, \gamma_v$ | Function variances (development, inheritance) |
| $D(z' \vert z, N, t)$ | Inheritance function |
| $E(a_m, K)$ | Life expectancy at sexual maturity and carrying capacity |
| $G(z' \vert z, N, t)$ | Development function |
| $K$ | Carrying capacity |
| $\mathbf{K}$ | Vector of population size structure at $K$ |
| $L(a, K)$ | Survivorship to age $a$ evaluated at $K$ |
| $L(a_m, K)$ | Proportion of each cohort surviving to $z_m$ evaluated at $K$ |
| $\lambda$ | Population growth rate |
| $N$ | Population size |
| $N(z, t)$ | Distribution of body size at time $t$ |
| $p_a$ | Proportion of the population that is sexually mature |
| $p_{a,K}$ | Proportion of the population that is sexually mature at $K$ |
| $p_j$ | Proportion of the population in the juvenile age class |
| $p_{j,K}$ | Proportion of the population in the juvenile age class at $K$ |
| $R_0$ | Mean lifetime reproductive success |
| $R(z, N, t)$ | Reproduction function |
| $R(a, K)$ | Per capita reproductive success at age $a$ evaluated at $K$ |
| $\bar{R}_{a,K}$ | Mean adult reproductive rate at $K$ |
| $\theta(\mu = \dots, \sigma^2 = \dots)$ | Normal distribution with mean $\mu$ and variance $\sigma^2$ |
| $S(z, N, t)$ | Survival function |
| $\bar{S}_{x,K}$ | Mean adult survival at $K$, with $x = j$ for juveniles and $x = a$ for adults |
| $t$ | Time |
| $z$ | Body size |
| $z_m$ | Size at sexual maturity |
| $\tilde{z}$ | Body size at which mean juvenile survival is observed |
| $\bar{z}_j$ | Mean body size of juveniles |

### The model

We use a class of model called an integral projection model (IPM) (Coulson, 2012; Ellner et al., 2016). Each unique parameterization of an IPM describes a life history strategy (Childs et al., 2004; Kentie et al., 2020; Metcalf et al., 2008), and each IPM projects population dynamics of that strategy (Ellner et al., 2016). These attributes make IPMs ideally suited to explore life history evolution (Childs et al., 2004).

We develop a size-structured integral projection model (IPM) that consists of four equations describing the association between body size *z* at time *t* and i) survival to time 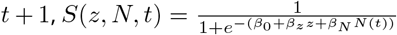, ii) the growth trajectory of surviving individuals from *t* to 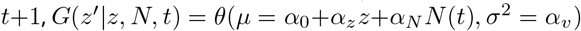, iii) the per-capita reproductive rate between *t* and *t*+1 defined as the number of offspring produced immediately after the population census at time t that survive to recruit to the population at time *t* + 1,

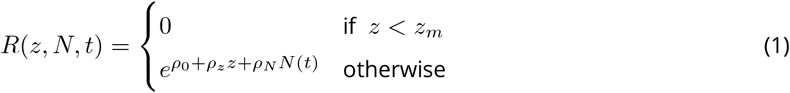

and iv) the body size of these offspring at recruitment to the population at 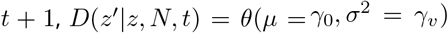 where the *θ*s are normal distributions with means of *μ* and variances *σ*^2^, the *α*s, *β*s, *γ*s and ρs are parameters, and *z_m_* is size at sexual maturity. These four functions combine to iterate forward the distribution of body size *N*(*z, t*) within the population at time *t* to the distribution of body size *N*(*z*’, *t* + 1) at time *t* + 1:

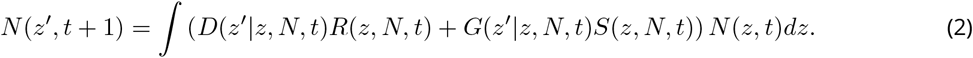

We assume a pre-breeding census such that reproduction captures the production of offspring and their survival from birth to recruitment to the population at t +1. When we refer to density-dependent reproduction, negative density dependence can impact either of these two processes as is standard in discrete density-dependent models with a pre-breeding census (Caswell, 2001; Charlesworth, 1994). The function *G*(*z*’|*z, N, t*) that describes growth trajectories is called the development function as is standard nomenclature in IPM notation (Coulson, Kendall, et al., 2017), and describes growth from one age to the next. IPMs can be constructed for any continuous phenotypic trait – not just body size – and the function can be mechanistic, capturing detailed developmental pathways, or phenomenological based on repeated phenotypic measurements taken on the same individuals over time (Ellner et al., 2016; Lachish et al., 2020; Smallegange et al., 2017).

Because this is a density-dependent model, at equilibrium *N*(*z, t*) = *N*(*z*’,*t* + 1). We discretise each of the functions to allow us to approximate the integral projection model in matrix form using standard approaches (Ellner et al., 2016). At equilibrium, the approximation is **K** = (**DR** + **GS**)**K** where **K** is a vector describing the population size structure at carrying capacity, and each emboldened letter represents a matrix capturing the similarly named function in equation 2.

In our models we set some slopes to zero to remove the effects of either body size or density dependence on either survival or reproduction. We do this to keep our models simple. By doing this, we only include density dependence in one function at a time. In the first scenario, density dependence acts on reproduction, limiting the number of offspring produced. We modeled this by setting *ρ_N_* < 0 (Equation 1). In the second scenario, population size is controlled via juvenile survival such that density has a negative effect on survival for juveniles but not for adults, and we modeled this via setting

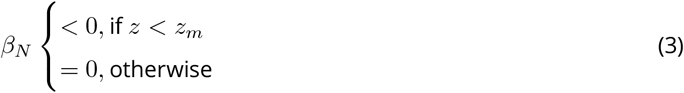

We refer to these two scenarios as “density-dependent reproduction” and “density-dependent juvenile survival” respectively.

At equilibrium, when the population size of a life history is at carrying capacity, both the population growth rate λ and mean lifetime reproductive *R*0 are equal to one and the dominant eigenvalue of the matrix approximation is 1 (Caswell, 2001).

### Iterating the model

Our analysis proceeds by iterating a population with a given life history strategy through time until it reaches a constant population size, *K* (Coulson, 2012; Ellner et al., 2016). Because these models are ergodic, the same equilibrium is achieved independent of the initial population size structure. We consequently generate a random population structure at time *t* = 1 and then numerically iterate the population forward until a stable population size and size structure is achieved. At each iteration we use population size at time *t* to determine the values of the density-dependent function used to project the population forward from *t* to *t* + 1. We then report quantities such as the proportion of the population that is sexually mature, life expectancies at a given age, and the probability of achieving sexual maturity at K for each life history strategy. We use these quantities to identify circumstances when extremes of body size and life history evolve.

### Defining life history strategies

Within each of the two scenarios, we construct 20 models, each representing a different life history strategy with different growth trajectories and sizes at sexual maturity. Within a scenario, each of these 20 models has identical parameter valuesforeach function, with the exception of the development function *G*(*z*’|*z, N, t*) and the size at sexual maturity *z_m_* which is always 80% of asymptotic size, which means that *z_m_* is an emergent property of the development function specific to each life history strategy. Different parameterisations of the development function generate different stable size distributions (the dominant right eigenvector of the IPM evaluated at *K*) for each life history, and these differences generate variation in age-specific survival rates (see results). Demographic rates must combine to give λ = *R*0 = 1 at equilibrium. Because survival rates vary across life history in both scenarios, the one degree of freedom available within the model to satisfy the condition λ = *R*0 = 1 at equilibrium will be the value of *K* in the density-dependent function (reproduction or juvenile survival). For each model, we find the value of *K* via numerical iteration (see above). The life history with the largest value of *K* will be the fittest, and in an evolutionary game would always grow to dominate the population if we assume that individuals are competitively equivalent across strategies - i.e. symmetric competition.

We keep the models simple by assuming that each reproducing parent produces the same distribution of offspring body sizes regardless of their size or life history strategy (Fig 1(A)). Body size is consequently not heritable within each life history strategy (Plard et al., 2021), but each life history strategy is passed from generation to generation with perfect fidelity (Childs et al., 2004). We also assume that all offspring initially develop at the same pace regardless of life history strategy. After age 1, the development functions diverge among the life histories (Fig 1(B)), such that those that will go on to achieve a larger size and greater age at sexual maturity continue to develop quickly, while those that will mature at a smaller size and lesser age slow their growth rates, reaching their asymptotic sizes at a younger age (Fig 1(C)). The growth models are monomolecular, such that growth rate slows with increasing size. We choose this formulation because monomolecular growth (i) is a good descriptor of growth in many species, and (ii) can be described with fewer parameters than non-linear growth forms (English et al., 2012; Gaillard et al., 1997).

**Figure 1.**
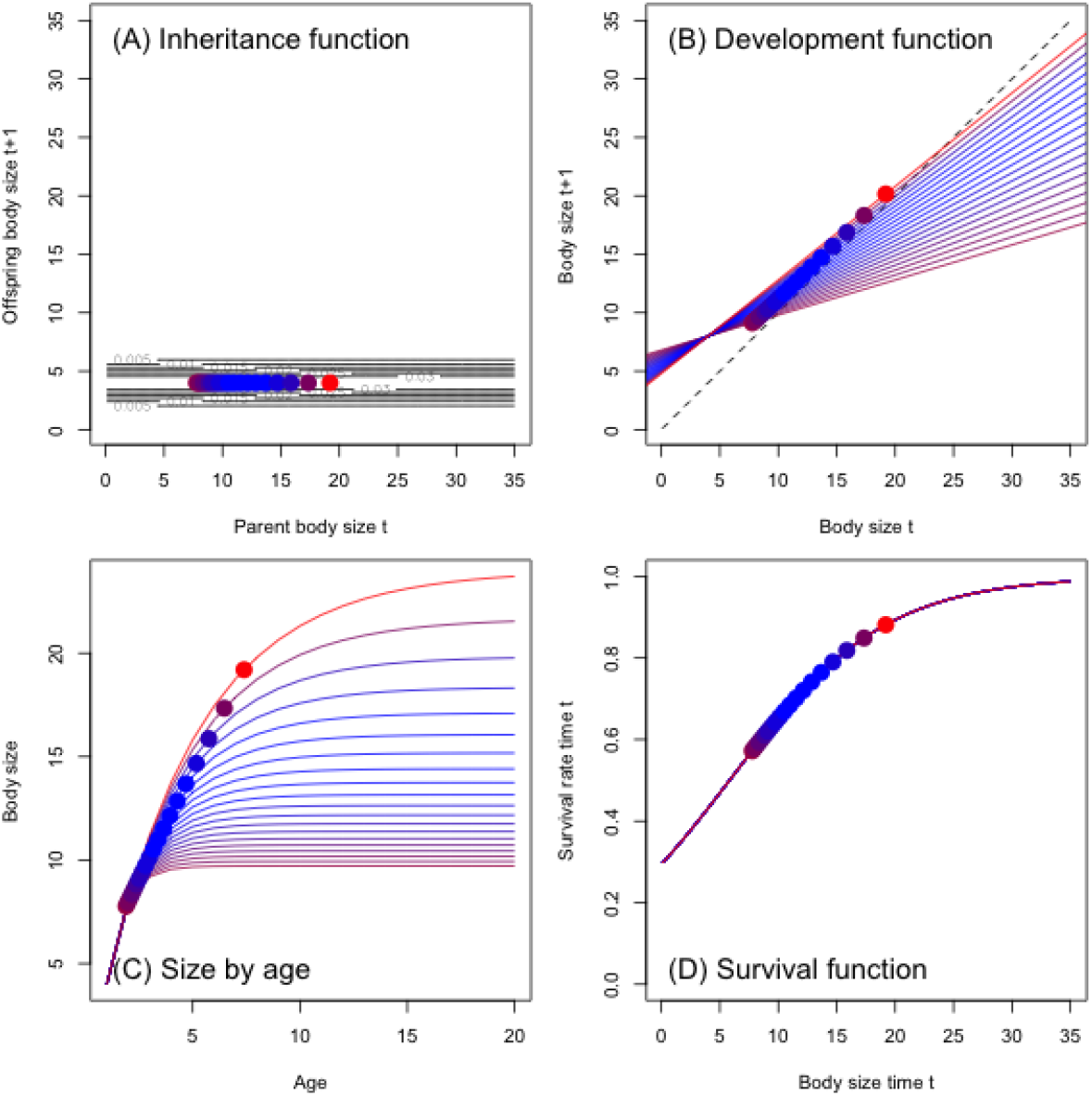
Density-independent functions in the density-dependent reproduction scenario. (A) association between parental size at time *t* and offspring size at time *t* + 1, where the black lines are contours representing the probability distributions of offspring sizes, (B) development functions, where contours representing the probability distribution of sizes at t+1 are not shown, (C) monomolecular growth functions showing size as a function of age, (D) mean of the body size-survival function. Unlike in A) we do not show the contours of the probability distributions. Each line represents one of the 20 life histories, and each dot represents size (or age in the case of C) at sexual maturity. The redder the colour of a point, the fitter the life history strategy (i.e., greater carrying capacity).

Survival rates increase with body size in all life histories in the same manner (Fig 1(D)), although when density dependence operates on juvenile survival this function is depressed when *z* < *z_m_* for each life history. Reproduction does not vary with size, i.e. *ρ_z_* = 0 in both the density-dependent reproduction and density-dependent juvenile survival scenarios, but the elevation of the function does vary with population density in the scenario where reproduction is density-dependent. Parameter values (Table 2) differ between the two scenarios to enable us to more easily graphically depict dynamics.

**Table 2.** Parameter values used in the two scenarios.

| Function | Intercept | Body size slope | Density slope | Variance intercept |
| --- | --- | --- | --- | --- |
| Density-dependent reproduction - scenario 1 |  |  |  |  |
| Survival | $\beta_0 = -0.875$ | $\beta_z = 0.15$ | $\beta_N = 0$ | |
| Reproduction | $\rho_0 = 1$ | $\rho_z = 0$ | $\rho_N = -0.001$ | |
| Density-dependent juvenile survival - scenario 2 |  |  |  |  |
| Survival | $\beta_0 = 0.25$ | $\beta_z = 0.125$ | $z < z_m : \beta_N = -0.001$ else 0 | |
| Reproduction | $\rho_0 = -1$ | $\rho_z = 0$ | $\rho_N = 0$ | |
| Both scenarios |  |  |  |  |
| Growth life history 1 | $\alpha_0 = 6.8$ | $\alpha_z = 0.3$ | $\alpha_N = 0$ | $\alpha_v = 1$ |
| Growth life history 2 | $\alpha_0 = 6.69$ | $\alpha_z = 0.33$ | $\alpha_N = 0$ | $\alpha_v = 1$ |
| Growth life history 3 | $\alpha_0 = 6.59$ | $\alpha_z = 0.35$ | $\alpha_N = 0$ | $\alpha_v = 1$ |
| ... | ... |  |  |  |
| Growth life history 20 | $\alpha_0 = 4.8$ | $\alpha_z = 0.8$ | $\alpha_N = 0$ | $\alpha_v = 1$ |
| Inheritance | $\gamma_0 = 4$ | NA | NA | $\gamma_v = 1$ |

### Interpreting model outputs

We start by examining the association between size at sexual maturity *z_m_* and carrying capacity *K* to characterize selection on life history strategy. We then wish to biologically and mathematically explain why the patterns we observe are generated.

Our first objective is to gain insight into how carrying capacity is maximized. We calculate terms describing the population dynamics and examine how these vary with size at sexual maturity across life history strategies within each scenario. Those terms that show similar associations to those we identify between size at sexual maturity and carrying capacity must be major drivers of the dynamics.

To do this, we start by writing the population dynamics as a function of mean class-specific demographic rates. Because the model distinguishes juveniles *z* < *z_m_* and adults *z* ≥ *z_m_*, it helps to write the population dynamics as a function of juvenile and adult rates. Specifically, we write the population growth rate, λ = 1 at equilibrium as a function of the proportion of juvenile *p_j_* and adult age classes (*p_a_* = 1 – *p_j_*) in the population and their mean survival (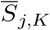 and 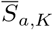) and reproductive rates (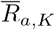 remembering juveniles do not reproduce),

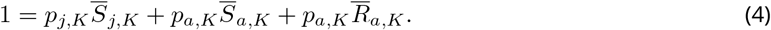

We next rearrange equation 4 to put the density-dependent rate on the left hand side. We also drop the subscript *K* for the density-independent rates. Next, we replace the mean value of the density-dependent demographic rate with the equation that describes the rate. For example, recall that 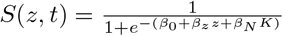. Mean survival in the density-dependent juvenile survival scenario is 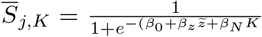, where 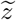 is the value of *z* that produces the mean survival rate across the distribution of juvenile body sizes. Note that nonlinear averaging means that 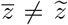. Finally, we rearrange and simplify the resulting equation to have K on the left hand side. The density-dependent reproduction and density-dependent juvenile survival scenarios respectively produced the following expressions

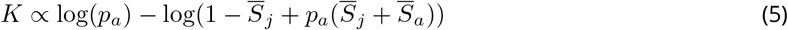

and

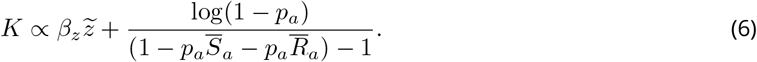

Through this rearrangement, we now have functions describing fitness (i.e. *K*) for each scenario. We calculate each of the terms in these expressions using approaches in Coulson, Tuljapurkar, et al. (2010) and then examine how each term is associated with size at sexual maturity across life history strategies within each scenario.

Having identified the factor that determines carrying capacity in both scenarios, the logical next step was to explore how evolution optimizes survivorship and fertility schedules. To do this, we write the life histories as a function of survivorship and fertility schedules. Because our developmental functions are continuous, we choose to write these schedules in continuous time, but they could easily be written as summations instead of integrals. For an age-structured density-dependent life history at carrying capacity we can write the Euler-Lotka identity as

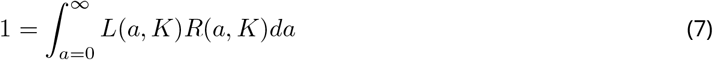

where *L*(*a, K*) and *R*(*a, K*) are respectively survivorship to age *a* and per-capita reproductive success at age a, both evaluated at carrying capacity, *K*. Because reproduction does not occur until sexual maturity is reached

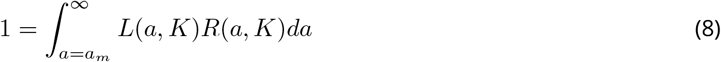

where *a_m_* is age at sexual maturity. In the models *R*(*a, K*) is constant across ages beyond sexual maturity within a life history so we simplify to *R*(*K*) then write,

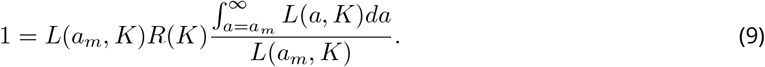

The survivorship term *L*(*a_m_, K*) is the proportion of each cohort surviving to sexual maturity, and 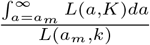 is life expectancy at sexual maturity that we write as *E*(*a_m_, K*). This reveals a trade-off between per-capita reproduction, the proportion of each cohort surviving to sexual maturity, and life expectancy at sexual maturity. In the density-dependent reproduction scenario, *R*(*a,K*) is density-dependent, so we separate the density-dependent and -independent rates, such that,

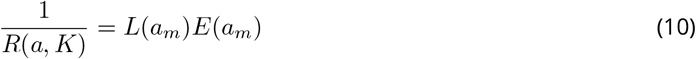

and

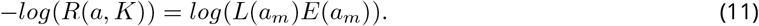

We therefore expect to see a negative linear association depicting a trade-off between the product of life history traits that are density-independent, with the value of density-dependent life history traits. We use an identical approach for the density-dependent juvenile survival scenario.

We calculate these continuous age-structured quantities by using Steiner et al. (2012)‘s derivation of a stage duration matrix, **P** = (**I** – **T**)^-1^ where **I** is the identity matrix and **T** = **GS**. Each *i,j* element in this matrix describes the expected amount of time an individual in stage i will spend in stage *j* before death. We can sum these elements across columns to calculate life expectancy for an individual at sexual maturity, and across rows to calculate survivorship from birth to the size at sexual maturity (Steiner et al., 2012).

## Results

### Disruptive selection on body size

In both the density-dependent reproduction and density-dependent juvenile survival scenarios we observe disruptive selection on body size (Fig 2(A,B)). Below a threshold size at sexual maturity where lowest carrying capacity is observed there is directional selection for small size at sexual maturity and a fast life history. Above the threshold, evolution of gigantism is observed. Why do we observe these patterns?

**Figure 2.**
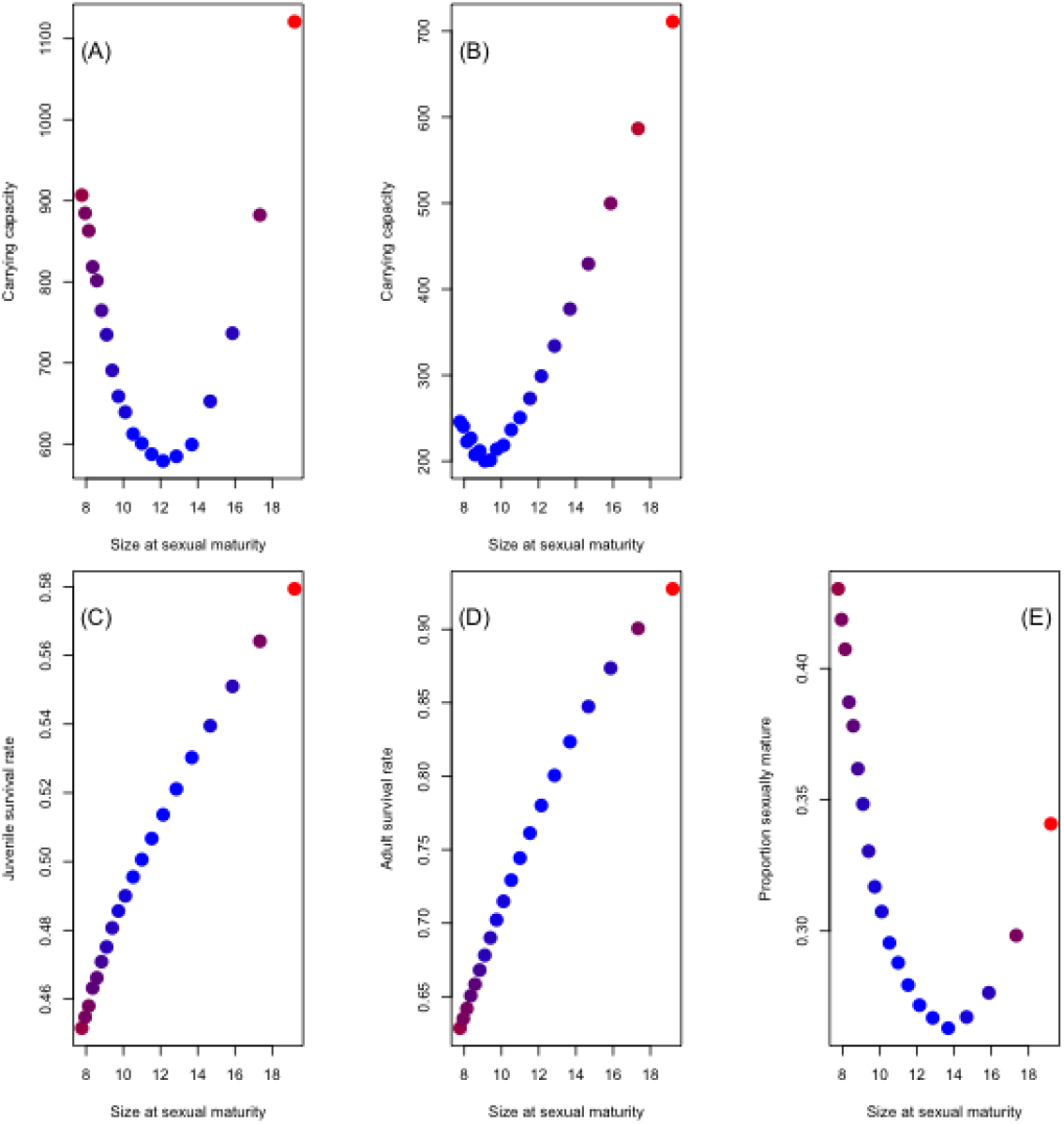
Scenario outcomes and why different life histories have different carrying capacities at equilibrium. Disruptive selection on size at sexual maturity for (A) the density-dependent reproduction scenario and (B) the density-dependent juveniles survival scenario. Each point represents one of our 20 life histories. The redder the colour of a point, the fitter the life history strategy. Association between size at sexual maturity and (C) juvenile survival, (D) adult survival, and (E) proportion of the population that is sexually mature for the density-dependent reproduction scenario.

### Maximizing carrying capacity

Because carrying capacity is fitness, as it increases across life history strategies, the predicted value of density-dependent terms in models will decrease. For example, in the density-dependent reproduction scenario (equation 1), the strategy with the highest carrying capacity will have the most negative value of the term *ρ_N_K*, and, on the scale of response, the smallest value of 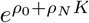. In the density-dependent juvenile survival scenario, the strategy with the highest carrying capacity will have the most negative value of the term *β_N_K* and, on the scale of response, the smallest value of 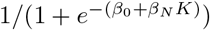.

### Factors determining carrying capacity

In the density-dependent reproduction scenario we find that of the three terms in equation 5, mean juvenile (Fig 2(C)) and mean adult survival (Fig 2(D)) are positively associated with size at sexual maturity, while the proportion of the population that is sexually mature (Fig 2(E)) exhibits a “u”-shaped relationship of similar form, but with a different minimum, to the pattern of disruptive selection seen in Fig 2(A).

We find a similar pattern in the density-dependent juvenile survival scenario using equation 6 with the proportion of the population that is sexually mature exhibiting a “u”-shaped association with size at sexual maturity and the other terms exhibiting positive associations (Fig S1). These results suggest that understanding the dynamics of the proportion of the population that is sexually mature is central to understanding the patterns we observe, and that requires understanding survivorship and fertility schedules. We consequently now turn our attention to the dynamics of life histories.

### Life history dynamics

We start by considering the density-dependent reproduction scenario. Holding the size-survival function constant (Fig 1(D)), but altering the development function (Fig 1(B,C)), inevitably changes the survivorship function: the probability of surviving from birth to any given age (Fig 3(A)). The faster that individuals grow, the more quickly they progress along the x-axis of the body size-survival function (Fig 1(D)), and this means that their probability of surviving to, and at, each age increases when going from fast-lived to slow-lived life histories.

**Figure 3.**
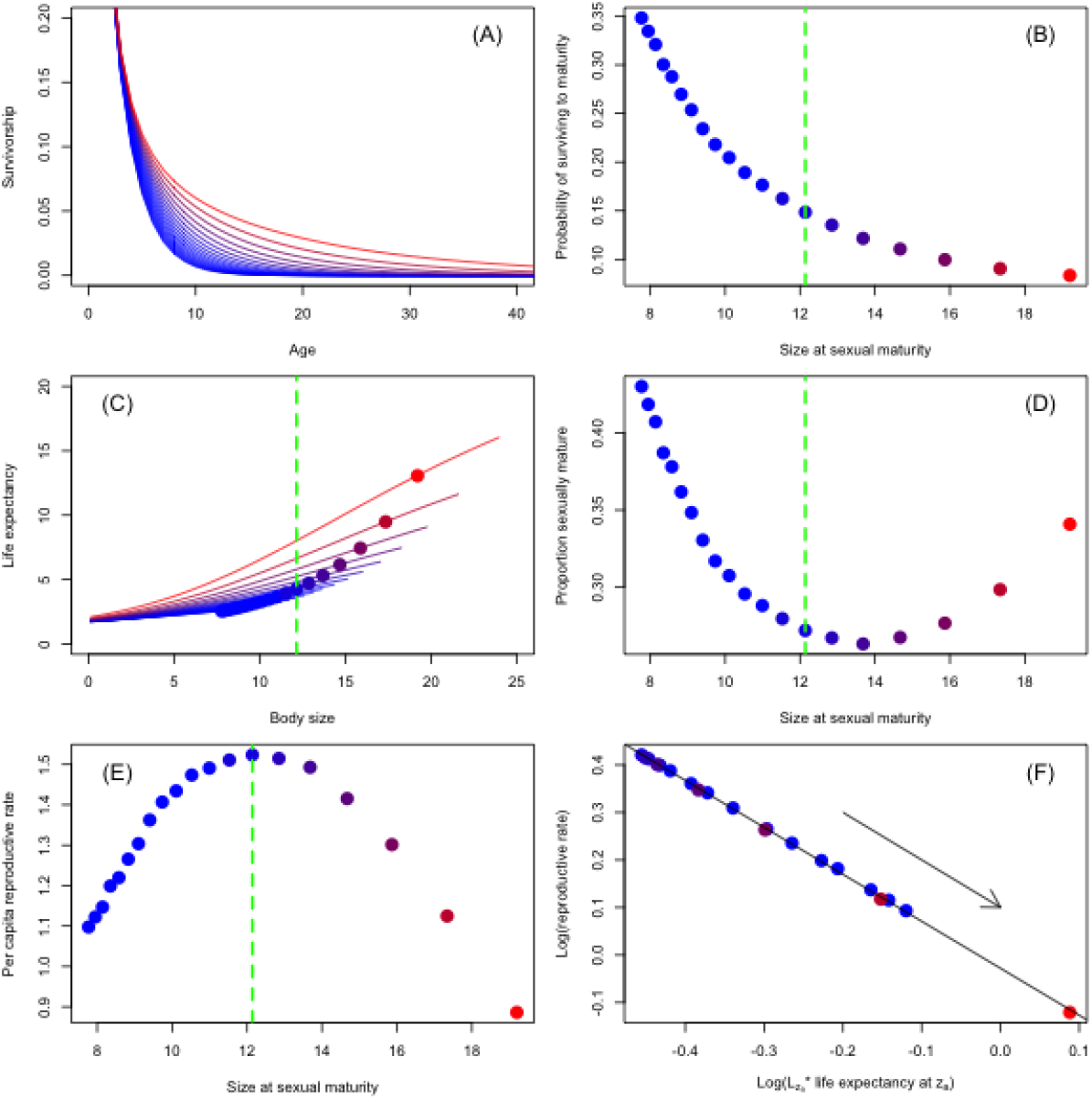
Figure 3. Model dynamics for the density-dependent reproduction scenario. (A) survivorship functions for each life history, (B) survivorship to sexual maturity as a function of size at sexual maturity, (C) life expectancy as a function of body size, (D) proportion of population that is sexually mature as a function of size at sexual maturity, and (E) per-time step per-capita reproductive rate as a function of size at sexual maturity, and (F) trade-off between the log of the density-independent rates with the log of the per-capita per-time step reproductive rate (the arrow represents the direction of evolution). The x-axis label is the combination of the density-independent life history traits. The dotted green vertical lines in (B-E) represent the life history of minimum fitness (i.e., lowest carrying capacity). Fitness increases either side of the green line. Each point represents one of our 20 life histories. The redder the colour of a point, the fitter the life history strategy. Note that Fig 3(D) uses identical data to Fig 2(E) but in Fig 3(D) we draw the vertical line of minimum fitness.

The change in the development function, and in size and age at sexual maturity, generates variation in the probability of an individual surviving to sexual maturity across life histories (Fig 3(B)). A smaller proportion of each cohort achieves sexual maturity as size at sexual maturity increases because it takes longer to achieve sexual maturity, and this delay imposes a greater mortality burden on each cohort than the survival benefits accrued via achieving larger sizes at a particular juvenile age. The mortality cost of delaying sexual maturity can be offset by an increase in life expectancy at sexual maturity (Fig 3(C)) as larger adults have higher per-time step survival rates than those that are smaller (Fig 1(D)) and consequently live for longer.

Below the threshold of minimum fitness (green line in Fig 3(B-E)) the proportion of the population achieving sexual maturity decreases at a relatively faster rate than the corresponding increase in life expectancy, with the converse true above the threshold. A consequence of these contrasting rates of change is that the proportion of sexually mature individuals within the population can increase (Fig 3(D)), even though a smaller proportion of each cohort achieves sexual maturity (Fig 3(B)), simply because a greater number of cohorts are alive as adults at any one time as adult life expectancy increases. Once individuals achieve sexual maturity, they reproduce.

The switch in the relative sizes of the derivative of the proportion of each cohortsurvivingto sexual maturity to size at sexual maturity, and the derivative of life expectancy to size at sexual maturity, generates disruptive selection. We observe an “n”-shaped association between size at sexual maturity and the per-capita reproductive rate (Fig 3(E)), which is reflected in a mirror-image “u”-shaped association between size at sexual maturity and carrying capacity (Fig 2(A)). The constraint *R*0 = 1 means that the minimization of the density-dependent term in the density-dependent reproduction function must be countered by maximization of values predicted by the density-independent body size term in the survivorship function (*β_z_z*). Because the survivorship function determines both the proportion of each cohort that achieves sexual maturity, and life expectancy at sexual maturity, and given equation 11, we observe a linear association with a slope of –1 between the log of the product of survivorship to sexual maturity and life expectancy at sexual maturity with the log of the per-capita reproductive rate (Fig 3(F)).

In our second scenario, where juvenile survival is density-dependent, survival is dependent on body size as well as population size. Reproduction is now density-independent and *ρ_N_* = 0. A consequence of these changes is the form of the survival and survivorship functions now differ compared with the density-dependent reproduction scenario. The density-independent terms are now the effects of body size on juvenile and adult survival (*β_z_z*), while per-capita reproduction does not vary with life history because *ρ_z_* =0 and *ρ_N_* = 0.

As before, the probability of surviving to maturity declines with increasing size at sexual maturity, while life expectancy increases. These processes combine to generate a quadratic association between size at sexual maturity and the proportion of the population that is sexually mature. The same maximization of K, and minimization of the density-dependent term occurs as in the density-dependent reproduction scenario, except the demographic rate that is modified is *S*(*z* < *z_m_, N, t*), and the term being minimized is now *β*_0_ + *β_N_K*. The density-independent life history quantity that is maximized is adult life expectancy.

There is one significant difference between the two scenarios: survival, unlike reproduction, is a function of body size. Because the development function varies across life histories along with size at sexual maturity, mean juvenile body size, and mean juvenile survival, also vary with life history. A consequence of the role of body size on juvenile survival is that the life history with minimum fitness does not align with the life history that has the maximum per-capita juvenile survival rate. This does not affect the negative linear association between the logs of the density-independent and density-dependent rates. Figure S2 provides an equivalent version of Figure 3 for the density-dependent juvenile survival scenario.

We can now understand why disruptive selection is driven via the proportion of the population achieving sexual maturity observed in our analyses of *K*. There is a trade-off between the mortality rates experienced by juveniles and the survival and reproductive rates of the sexually mature. The trade-off is mediated by rates of development. Size at sexual maturity is selected to increase when the fitness benefits for sexually mature adults of achieving a large size by delaying the age of sexual maturity outweigh the mortality costs to juveniles caused by delaying the age of sexual maturity. When this occurs, we see selection for gigantism and slow life histories. In contrast, size at sexual maturity is selected to decrease when the fitness benefits to the sexually mature are less than the mortality cost endured by juveniles. The point at which the trade-off switches, generating disruptive selection, is dependent upon where in the life history density dependence operates.

In Figure 4 we schematically illustrate this dynamic. The summary figure does not include body size because its inclusion complicates visual interpretation. The figure shows how a change in age at sexual maturity (4(A) versus 4(B)) results in a change in the form of the survivorship function, which results in a change in the elevation of the density-dependent reproductive function to ensure *R*_0_ = 1. The life history in Figure 4(B) is favoured by selection in this example because the density-dependent reproductive function is ata lower elevation than in Figure 4(A). Figure 4(C) provides an explanation of the rectangle approximation used in equation 9.

**Figure 4.**
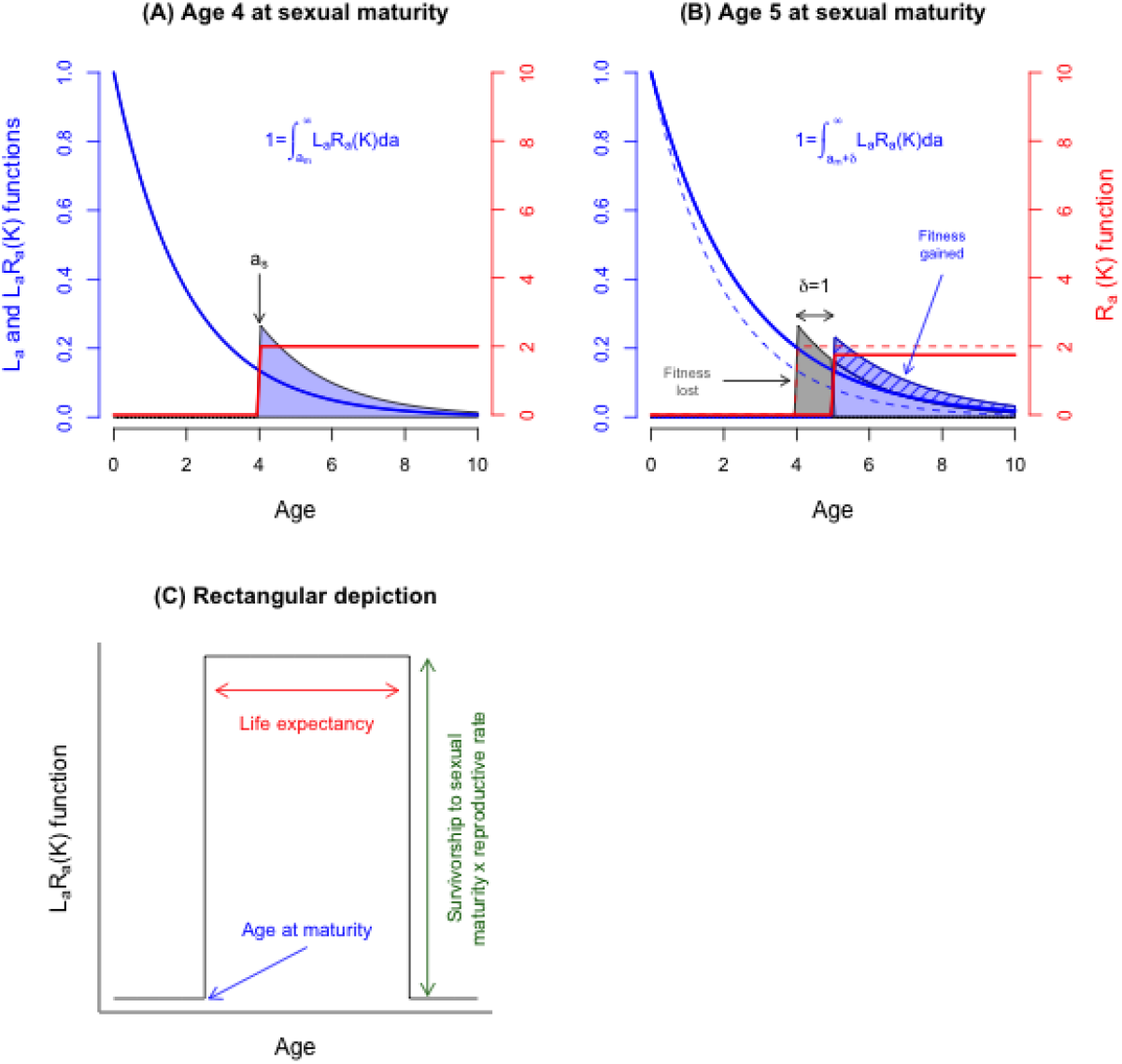
Summary of the age-structured life history dynamics of the model where reproduction is density-dependent. Blue lines represent the survivorship schedules (left y-axis) and red lines the fertility schedules (righty-axis). The polygons represent the product of the survivorship and reproduction schedules that are described with the equations. The initial life history strategy is depicted in (A), the mutant strategy, with a delayed age at sexual maturity, in (B). The delay in age at sexual maturity results in a change in the development function that results in an elevation of the survivorship function (compare the solid blue line in (B) to the solid blue line in (A) which is also represented by the dotted blue line in (B)). Because the volume of the blue polygon in (A) and (B) must equal one (equations on plot), the reproduction function is depressed in (B) compared to (A) (compare the solid red lines in (A) and (B)). The grey and hashed blue polygons in (B) show how the polygon has changed shape between the two life histories. (C) Rectangular approximation of the life history function used to identify trade-offs.

These results suggest that if we alter the fitness costs and benefits of delaying sexual maturity, we should be able to shift the size at sexual maturity at which we see a switch in the direction of selection. We explore this by modifying the survival function in the density-dependent reproduction scenario.

### Changing the size-survival function

The rate at which survival changes with age determines why the proportion of each cohort that achieves sexual maturity changes at a different rate across the life histories than life expectancy at sexual maturity. The elevation and slope of the size-survival function should consequently determine selection on life history. We examined this for the density-dependent reproduction scenario by systematically modifying the intercept and slope of the survival function *S*(*z, N, t*) (Fig 5).

**Figure 5.**
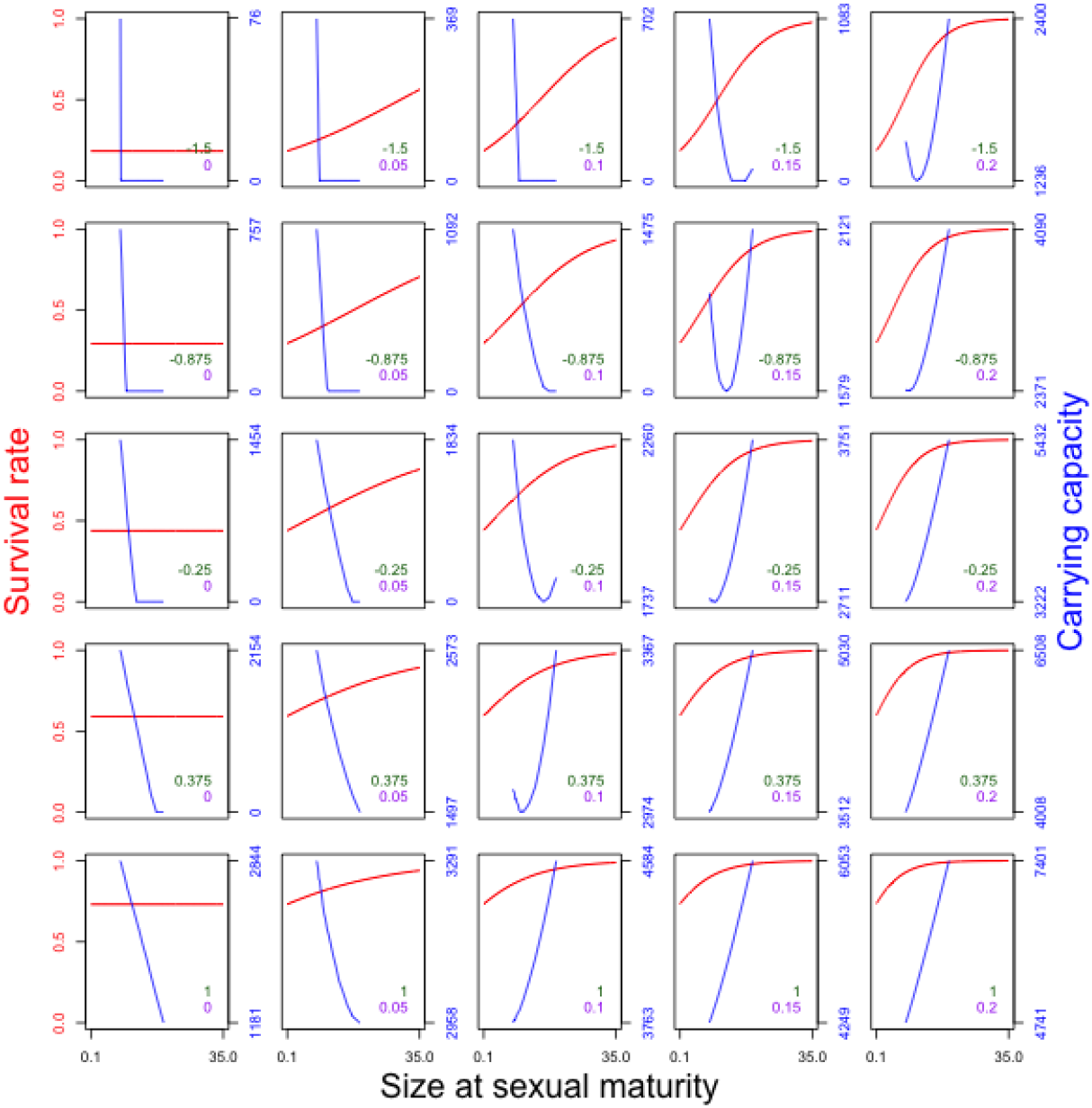
Figure 5. Dynamical consequences of altering the intercept and slope of the body size-survival function in the density-dependent reproduction scenario. As the elevation of the intercept (rows, and green numbers) and steepness of slope (columns and purple numbers) are altered, the change in the size-survival function alters selection on size at sexual maturity. The red lines represent the form of the size-survival function. The blue lines show how carrying capacity changes across the range of sizes of sexual maturity.

When the slope of the body-size survival function is 0 we never observe selection for delayed age and size at sexual maturity and a slower life history (column 1). In order to see selection for an increase in size at sexual maturity, survival rates need to increase with body size (positive viability selection) and need to be sufficiently high for sexually mature adults to extend lifespan sufficiently to offset the costs of a smaller proportion of offspring surviving to sexual maturity (see equation 9). It is this fitness differential across ages that determines whether there will be selection for an increase or decrease in body size and age at sexual maturity.

Finally, to demonstrate that our results are not due to non-linearities in our model, we linearly approximated the model and explored outputs (Appendix). This revealed that the patterns we report are not a consequence of the linearities in our model functions.

The blue lines are analogous to Figure 2 for each parameterisation of the size-survival function but are drawn as continuous lines rather than as dots.

## Discussion

### Phenotypic traits and life history evolution

A large body of empirical research has revealed that numerous drivers can influence survival and reproduction in wild populations of animals and plants (Burke and Nol, 2000; Gimenez et al., 2012; Gulland, 1995; Major and Kendal, 1996; Toigo and Gaillard, 2003). These drivers can be classified as i) individual attributes such as age, sex, and phenotypic traits, ii) biotic drivers such as the size and structure of populations of the focal and interacting species, and iii) abiotic drivers such as the weather. The biotic and abiotic drivers limit population growth and size while individual phenotypic traits and their developmental trajectories evolve to minimize these biotic and abiotic impacts. Multiple phenotypic traits may be associated with a single limiting factor. By working within a framework where carrying capacity has repeatedly been shown to be fitness (Charlesworth, 1973,1994; Kentie et al., 2020; Lande, Engen, and Sæther, 2017; Lande, Engen, and Saether, 2009; MacArthur, 1962; Mylius and Diekmann, 1995; Takada and Nakajima, 1992, 1998), we reveal how evolution optimizes growth trajectories, survivorship, and fertility schedules that define the life history strategy. Optimization acts by minimizing the impact of population size on the density-dependent demographic rate. We find that when the adult fitness benefits of delaying sexual maturity to a greater age outweigh the juvenile mortality costs of doing so, gigantism can evolve. At the other extreme, small sizes at sexual maturity that are similar to those of Darwinian demons are selected. The key parameter driving these dynamics is the proportion of the population that is sexually mature, which is determined by the relative life expectancies of juveniles and adults.

Although our models are kept deliberately simple, they reveal important, general insights. First, the evolutionary stable life history strategy will always be the one that can persist at the highest impact of the limiting factors. In our models, the limiting factor is density. Density dependence is a dynamic that can be caused by various processes including predation and food limitation (Turchin, 1999). In a predator-limited environment, the evolutionarily stable life history strategy will therefore be the one that can persist at the highest predator density, while in a food limited case it will be the one that can either persist on the least available food or acquire a disproportionate amount of the food that is available. Thus how one dies, or how one is negatively affected by a density-dependent factor, impacts body size and life history evolution. Minimization of the impact of a limiting factor on the demographic rates it affects generates selection on phenotypic traits associated with surviving and reproducing in the factor’s presence (Coulson, 2021). If predation is the limiting factor, then camouflage or the ability to out-run a predator might be selected, while in a food-limited environment, traits subject to selection might be the ability to efficiently use energy, to migrate to greener pastures, or to defend a food source against conspecifics (Travis et al., 2014). Some of these traits change with age such that their dynamics are determined by developmental trajectories. When this is the case, these developmental trajectories are selected to optimize survivorship and fertility schedules to maximize fitness.

In some cases, there may be multiple factors that can limit a population via causing death or a failure to reproduce (Seip, 1992). For example, food shortages and pathogens may both contribute to limit a population. Different phenotypic traits and their developmental trajectories may be associated with each factor that causes death or a failure to reproduce. We have already described a couple of phenotypic traits that might be associated with food limitation; pathogens might drive selection on social behaviour and aspects of immune response.

In the presence of multiple causes of death or reproductive failure, evolution will optimize the development of many phenotypic traits simultaneously to determine the optimum age-specific survivorship and fertility schedules. When resources are limiting, being either hard to detect or acquire, this will generate trade-offs in their allocation (B Kooijman and S Kooijman, 2009). The fittest combination of traits will be the one that improves resource detection and acquisition while optimizing the allocation of resources to traits in a way that maximally reduces the likelihood of death or failure to reproduce from the limiting factors (Coulson, 2021). Despite all this complexity, if fitness can be defined for a particular environment, then the trade-off between the juvenile costs of delaying age at sexual maturity and the adult benefits of doing so will be general. The phenotypic details and energy budgets start to matter when mechanistic causes of the shape of survivorship and fertility schedules becomes the topic of interest (Lachish et al., 2020).

Two obvious questions arise from this conclusion: how do species of intermediate size and life history speed arise, and what about abiotic variation? Senescence is the decrease in survival or reproduction at older ages. We do not incorporate senescence into models, but given its ubiquitous nature (Nussey et al., 2013), it seems plausible that senescence means that survival and reproductive rates cannot remain indefinitely high among adults. Depending on the age at which senescence begins, and how quickly it happens, there could be a trade-off between rates of development, the shape of the survivorship and reproduction functions, and the onset of senescence (Jones et al., 2008). Future work should incorporate the effects of senescence into the framework we have developed to explore whether it can constrain the runaway selection our current models predict.

Abiotic variation can generate temporal variation in age- and trait-specific survival and reproductive rates and can also impact developmental trajectories (Tuljapurkar, 1990). Much in the same way that evolution will act to minimize the impact of a limiting factor on survival and reproduction, it will also select phenotypic traits to cope with abiotic variation (Lande, Engen, and Saether, 2009). In density-dependent stochastic environments where competition between individuals is symmetric, fitness is mean population size (Kentie et al., 2020). Depending upon circumstances that are not important for this discussion, an increase in abiotic variation can act to either increase or decrease mean population size (Tuljapurkar et al., 2009). If abiotic variation acts to decrease mean population size, then evolution will select for traits that improve individual resilience to abiotic variation, while if it increases mean population size, it will select for phenotypic traits that allow organisms to exploit the variation (Tuljapurkar et al., 2009). In future work we will develop this theme further.

A final avenue worth incorporating into models is the evolution of offspring size, which we kept constant in our models. Changing offspring size can also impact life history evolution (Charnov and Downhower, 1995; DA Reznick et al., 1990; Winkler and Wallin, 1987), and we can see two immediate impacts of altering the offspring number-offspring size trade-off. First, if carrying capacity is fitness, and density dependence operates via reproduction, then reducing litter size while increasing offspring size is one route to evolving a lower percapita reproductive rate allowing persistence at a higher carrying capacity (see also Parker and Begon (1986)). Second, larger offspring begin life further along the body size-survival function, potentially increasing the proportion of each cohort that survives to sexual maturity, altering the strength of selection on size at sexual maturity and life history speed. This second insight is novel and is only apparent after developing models like ours. Our framework will allow exploration of the evolution of offspring and litter size in a life history setting, and could be a valuable avenue of further research.

### Empirical considerations

Our work is theoretical, but it leads to a number of hypotheses that could be empirically tested. We show that the shapes of the four function types used to construct models determine whether small-bodied and fast, or large-bodied and slow, life histories are selected. To understand why a particular body size and life history evolves, it is consequently insightful to explore why the survival, development, reproduction, and inheritance functions take the shapes they do, and how they covary. What are the genetic, physiological, or environmental factors that determine the size-survival function, for example (Coulson, 2021)? As a population adapts to a new environment, the strength and form of feedbacks may change, and this will be reflected in the way the functions that constitute models change as adaptation occurs. Not only will this help us understand phenotypic trait and life history evolution, but also the way that the population dynamics change as adaptation occurs. We have examples from lab systems of how numerical dynamics changes with adaptive evolution or with different levels of genetic variation (Yoshida et al., 2003). Our approach offers ways to uncover mechanistic insight into what drives the co-evolution of traits and numerical dynamics as these are easily studied using IPMs (Coulson, MacNulty, et al., 2011). Understanding why we see particular functional forms, and how these change as adaptation progresses, will provide novel insight, but the approach also has the potential to help explain a number of evolutionary “rules”.

There are three main biogeograpical “rules” describing patterns of body size: the island rule, Bergmann’s rule, and Cope’s rule. The island rule states that small species of many mammals and birds tend to evolve large body sizes and slower life histories on islands, while larger species tend to evolve in the other direction (Clegg and Owens, 2002; Covas, 2012; Lomolino, 2005; Sandvig et al., 2019). Bergmann’s rule states that an increase in latitude typically corresponds to an increase in adult body sizes within species (McNab, 1971). Cope’s rule states that species tend to get larger over evolutionary time (Hone and MJ Benton, 2005), suggesting a similar process could well be happening over time as happens with latitude. These patterns suggest systematic changes in the shapes of size-survival, size-reproduction, development rates, and offspring size may underpin these “rules”. Additional work, where we impose fewer constraints on the functions in models, should help explain the circumstances required to generate these body size and life history patterns.

We can even hypothesize on the shape of the functions in extinct species, such as the giant sauropods. These giants are thought to have laid multiple clutches of relatively few ostrich egg-sized eggs, have very high early growth rates, and to achieve sexual maturity at around 30 years (Sander et al., 2011). The high growth rates suggested the young were unlikely food-limited, and selection for very large size suggests a steep increase in survival rates across the range of sizes through which they developed. Taken together, these suggest a high mortality rate on the young, likely via predation, but long life expectancies once sexual maturity was achieved.

## Conclusions

There are many ways in which the approach we use can be extended and models parameterized to address a range of empirical and theoretical questions about body size and life history evolution. In addition, our work also contributes to a general framework that we have been developing to study eco-evolutionary dynamics (Coulson, Kendall, et al., 2017; Coulson, MacNulty, et al., 2011). Our results reveal a general trade-off between juvenile and adult fitness that will determine age and size at sexual maturity and life history speed. They also help explain how a change in the predominant cause of death or failure to reproduce can result in predictable phenotypic trait and life history evolution in some species (D Reznick and Endler, 1982).

## Data accessibility

There is no data associated with this paper.

## Supplementary material

Scripts in R to run models at https://zenodo.org/record/7009178#.Yv9CaOwRXzc

## Acknowledgements

Thanks to Luke Coulson for running simulations over a range of parameter values. Thanks to Mike Furlong and the School of Biological Sciences at the University of Queensland for hosting TC’s and SC’s sabbaticals where the work was largely conducted. RDB is supported by NSF DEB2100163; Travis is supported by National Science Foundation award DEB 2100163; GP is funded by the UK’s Natural Environment Research Council through the Doctoral Training Partnership in Environmental Research at the University of Oxford (NE/L002612/1). Version 3 of this preprint has been peer-reviewed and recommended by Peer Community In Evolutionary Biology.

## Conflicts of interest disclosure

The authors of this preprint declare that they have no conflict of interest with the content of this article. Tim Coulson helped setup and is a recommender for PCI Ecology.

## Appendix

### Interpreting carrying capacity as fitness

Fitness is often considered to be genetic representation of a heritable entity (be it an allele, genotype, or strategy), either expected (Charlesworth, 1994) or realized (Coulson, T Benton, et al., 2006), in a population at some point in the future. Future genetic representation depends upon how quickly the heritable entity replicates and the degree of fidelity across generations (Fisher, 1930). Fitness is also often thought of as a growth rate, such as reproductive value (Grafen, 1999), or the speed at which an entity can invade a population of a resident (Dieckmann et al., 2006). When carrying capacity is fitness, it is shorthand for carrying capacity being the asymptotic endpoint of future representation of a heritable entity within a population at equilibrium, and whether one heritable entity would replace another in an evolutionary game (Kentie et al., 2020). For example, consider two competing strategies and assume that strategy *A* has a carrying capacity of *X* and strategy *B* of *X* – *q*. If one individual of strategy *B* were introduced into a population of strategy A at its carrying capacity *X*, it could not establish, because it would experience a population density that is greater than its carrying capacity. As a result, its replacement rate λ_*B*_, and its mean lifetime reproductive success R0B would both be less than one. In contrast, if the experiment were repeated the other way around, strategy A would have a growth rate λ_*A*_ > 1 and R0A > 1 because it would be introduced into a population below its own carrying capacity (Childs et al., 2004; Dieckmann et al., 2006; Meszena et al., 2002). If we know the carrying capacities of strategies *A* and *B*, we do not need to run an evolutionary game to identify the evolutionary endpoint (Kentie et al., 2020). Because carrying capacity is fitness in density-dependent environments, we can identify the evolutionarily stable strategy simply by finding the strategy with the largest carrying capacity.

### Linearisation of model

We linearised the model to demonstrate that the results are not a function of the non-linear aspect of our model.

We start with the simplification provided by equation 9 which we simplify the notation of to write 1 = *RJE* where *R* is reproduction, *J* juvenile survival, and *E* is life expectancy.

We can write 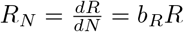 where *b_R_* is the density coefficient on an exponential *R*. If *P_a_* is survival at age a and *b_p_* the density coefficient on the logistic, then

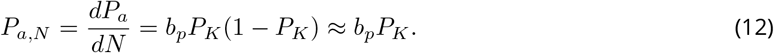

It follows that, approximately,

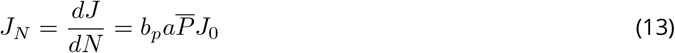

where 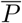 is average adult survival across the stable distribution of adult ages at *N*, and *J*_0_ is juvenile survival at *N* = 0.

Next, we make the density effect linear,

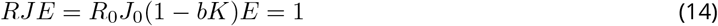

and

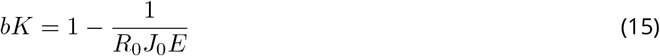

where *R*_0_ is reproduction evaluated at *N* = 0. Depending on the scenario, *b* = *b_R_* or *b* = *b_P_*.

For a range of *a* from *a_min_* to *a_max_* and a linear increase in survival rate with *P_a_* = *P*(*z_m_*) with a slope of *q*, then,

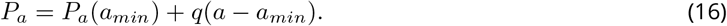

If we assume survival is constant post sexual maturity at *P_a_* then

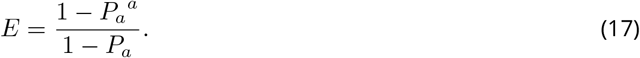

The slope of *E* now depends upon q as well as *a*, and, as in our simulations, life expectancy will only increase when *q* is large enough.

We can now use values of *E, R*_0_ and *J*_0_ to explore how linearised *K* varies as we change *E, b_P_* and *b_J_*. This is most easily done graphically. Mirroring our simulation results, divergent selection for *K* depends on a strong enough survival advantage of the delay in maturity. If not, *K* will just fall as *a* increases. Our results are consequently not due to the non-linearities in our functions.

**Figure S1.**
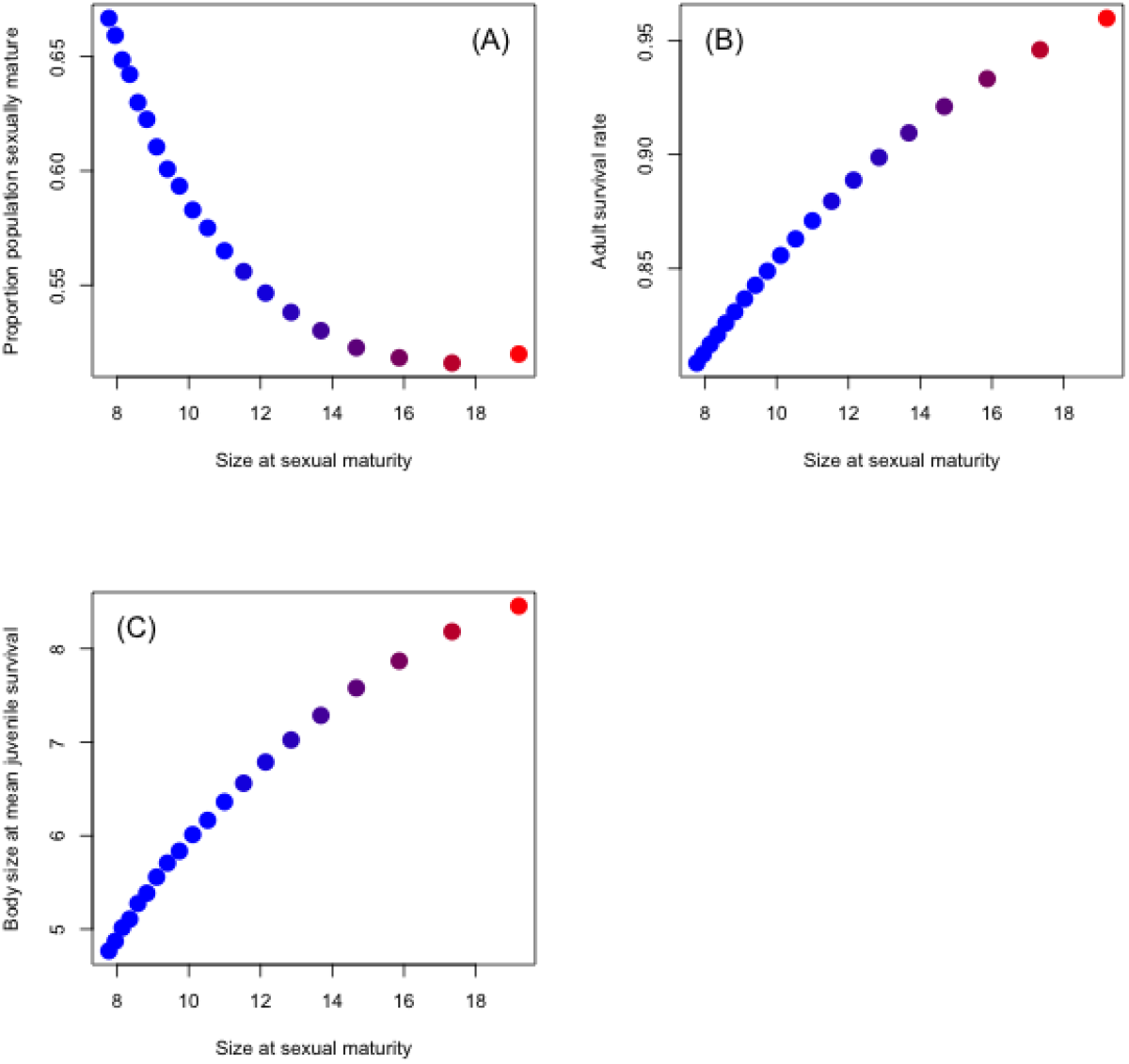
Associations between size at sexual maturity and each of the terms in our approximation of carrying capacity for the density-dependent juvenile survival scenario. Size at sexual maturity and (A) proportion of the population that is sexually mature, (B) adult survival and (C) body size of juveniles that predicts mean juvenile survival across the distribution of juvenile weights.

**Figure S2.**
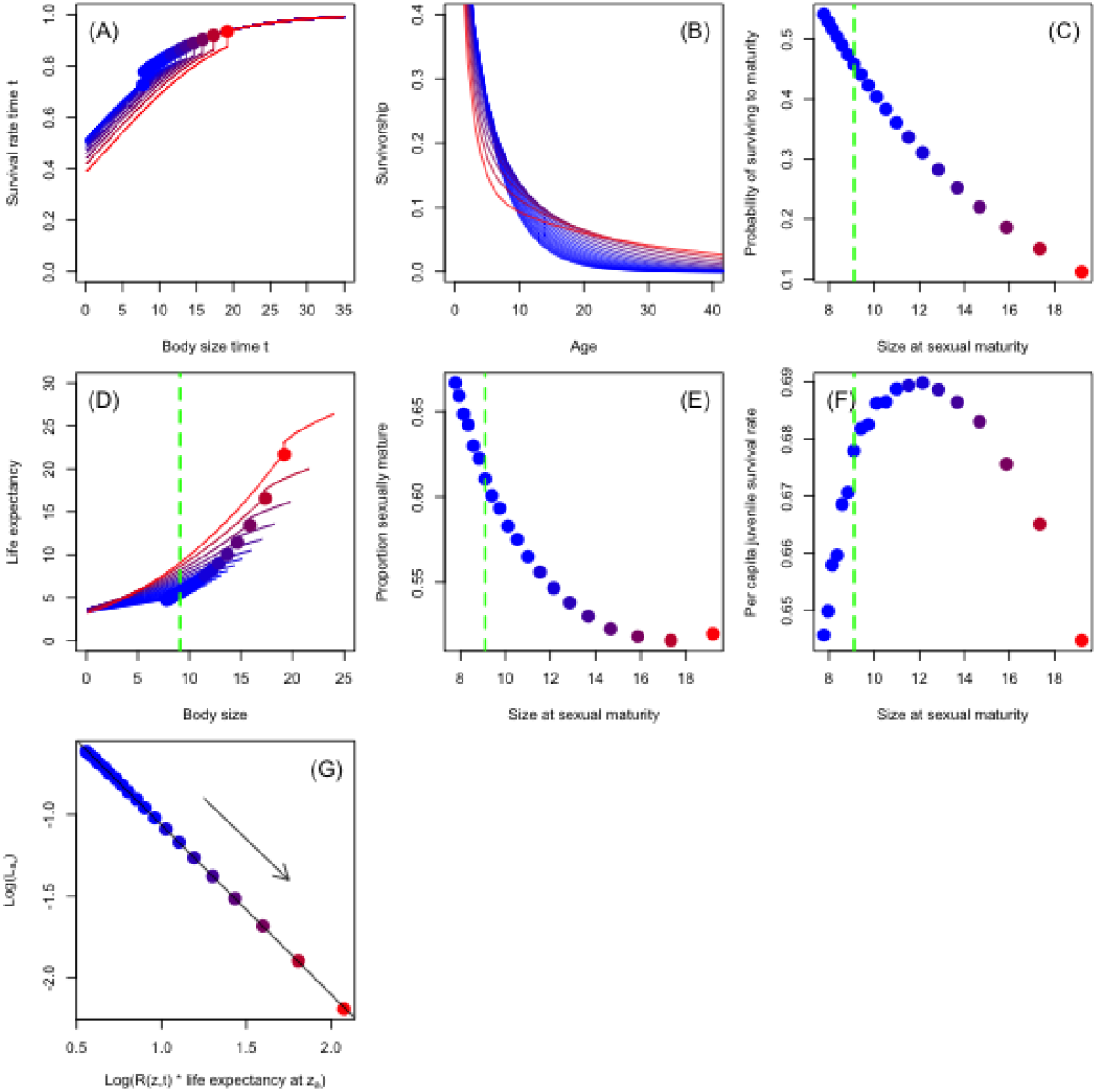
Model structure and outputs when the juvenile survival function is density-dependent (scenario 2). (A) body size-survival function, (B) survivorship functions for each life history, (C) survivorship to sexual maturity as a function of size at sexual maturity, (D) life expectancy as a function of body size, (E) proportion of population that is sexually mature as a function of size at sexual maturity, and (F) per-time step per-capita juvenile survival rate as a function of size at sexual maturity for each life history. (G) linear associations between the log of the density-independent rates against the log of the density-dependent rate (the arrow represents the direction of evolution). The dotted green vertical lines in (C-F) represent the life history of minimum fitness. Each point represents one of our 20 life histories. The redder the colour of a point, the fitter the life history strategy.

## Notes

https://zenodo.org/record/7009178#.Yv9CaOwRXzc

### Competing Interest Statement

The authors have declared no competing interest.

### Summary of Updates

Manuscript formatted to conform to PCI Ecology requirements following peer-review.

